# DIVERGENT TRAJECTORIES FOR ANAEROBIC MITOCHONDRIAL EVOLUTION IN BREVIATE PROTISTS

**DOI:** 10.64898/2026.01.29.702055

**Authors:** Zhenzhen Yi, Shelby K. Williams, Michelle M. Leger, Nikolaj Brask, Dayana Salas-Leiva, Jeffery Silberman, Yana Eglit, Alastair Simpson, Andrew J. Roger, Courtney W. Stairs

## Abstract

Mitochondrion-related organelles (MROs) have evolved as adaptations to low oxygen conditions multiple times in the eukaryote tree of life. However, the evolutionary steps by which aerobic mitochondrial functions were replaced by anaerobic pathways are still poorly understood. The breviate *Pygsuia biforma* is particularly interesting because it is the only protist known to have replaced the canonical mitochondrial iron-sulfur cluster (ISC) system with a horizontally acquired SUF-like minimal system (SMS) protein. This functions within an MRO possessing a uniquely configured electron transport chain (ETC). To investigate the evolutionary path by which the *P. biforma* MRO evolved these features, we conducted a comparative transcriptomic study of eight diverse marine breviate species and predicted their MRO proteomes.

We found three distinct patterns of iron-sulfur cluster biosynthesis machinery across the breviates where organisms would encode: the canonical ISC system alone, the ISC system and cytoplasmic SMS system, and a cytoplasmic and MRO-localized SMS system. Phylogenetic analyses suggests that the SMS system was acquired via lateral gene transfer in an ancestor of all breviates and later duplicated, with one copy gaining mitochondrial targeting and replacing the ISC system in a subset of breviates. We observed similarly divergent evolutionary trajectories for quinone-utilizing proteins. Two species have completely lost the ETC while the remaining lineages retain a partial ETC and mitochondrial contact site and cristae organizing system (MICOS), previously thought to be absent in breviates. These patterns reflect divergent biochemical configurations of MROs shaped by gene transfer, loss, and duplication within a single eukaryotic lineage and underscores dynamic remodeling of organellar metabolism in response to marine hypoxic environments.

## INTRODUCTION

Mitochondria are eukaryotic organelles that arose from endosymbiotic alphaproteobacteria prior to the last eukaryotic common ancestor (LECA)^1,2^. The mitochondria of most aerobic eukaryotes have an electron transport chain (ETC) that uses oxygen as a terminal electron acceptor, and generates ATP by oxidative phosphorylation^1^. However, some anaerobic eukaryotes possess so-called ‘mitochondrion-related organelles’ (MROs) that harbour metabolisms adapted to low-oxygen environments. For example, MROs have partially or completely lost their mtDNA and ETC, and instead rely on oxygen-independent pathways to biosynthesize ATP (reviewed in ^1,3–5^). The last decade of research into MROs from diverse protist lineages has demonstrated that these organelles perform a diverse spectrum of functions (reviewed in ^3,4^).

One highly conserved function of mitochondria and MROs is the biogenesis of iron-sulfur (Fe-S) clusters. Fe-S clusters are essential co-factors found in diverse organisms across the tree of life and are utilized in numerous proteins that are important for cellular metabolism, DNA replication, and redox biochemistry ^6^. In most eukaryotes, distinct systems for the maturation of Fe-S clusters occur in different subcellular compartments: an ‘iron sulfur cluster’ (ISC) pathway in mitochondria, a cytoplasmic iron-sulfur cluster assembly (CIA) system in the cytosol, and a sulfur mobilization (SUF) pathway in plastids ^7–9^. However, there are several deviations from this pattern. The ISC system has been completely lost, or replaced by other enzymes, in two unrelated lineages: a NIF-like system exists in the MROs^10,11^ or cytoplasm^12,13^ of some archamoebae, and an archaeal-like ‘SUF-like minimal system’ (SMS^14^) occurs in the MROs of *Pygsuia biforma* ^15^. In addition, an increasing number of anaerobic protist lineages have been shown to have a cytoplasmic SMS^16–18^ or bacterial-like SUF^19,20^ systems. Indeed, the acquisition of a cytosolic SUF system has been suggested to have preadapted some protists to the subsequent complete loss of the mitochondrial organelle^18,19^. Collectively, these findings demonstrate that the biosynthesis of Fe-S clusters varies across eukaryotes and has been shaped by lateral gene transfer and gene loss in various lineages.

Another highly conserved system of mitochondria and some MROs is the ETC. The ETC of aerobes typically consists of three proton pumping complexes (CI, CIII, CIV), one non-pumping complex (CII) and ATP synthase (CV). Some MROs have reduced variants of the ETC (reviewed in^21^); one frequently observed version is an ETC that retains CII but lacks CI, CIII, CIV and CV. In aerobic eukaryotes, CII oxidizes succinate to fumarate and electrons are transferred to ubiquinone (UQ) yielding ubiquinol. In some anaerobic eukaryotes, CII likely functions in reverse and uses an alternative lower-potential quinol called rhodoquinone (RQ), coupling fumarate reduction to the oxidation of rhodoquinol^22^. Although the source of reduced rhodoquinol is not known for MROs that lack CI, all protists that have only a CII always encode other quinone-reducing proteins in the ETC (e.g., electron-transferring flavoprotein dehydrogenase) that might be able to reduce rhodoquinone to rhodoquinol for CII-mediated fumarate reduction^21^.

Determining the evolutionary trajectories by which mitochondrial pathways were remodeled, including Fe-S cluster biosynthesis and respiration, is important for understanding how microbial eukaryotes become evolutionarily adapted to new environments, such as low-oxygen habitats. However, the ability to reconstruct these transitions has been limited by sparse sampling within major anaerobic lineages, leaving key steps in organellar evolution unresolved. Here, we addressed this knowledge gap by sequencing the transcriptomes of six diverse breviate protists and comparing their predicted MRO proteomes. Breviates (Breviatea) are small heterotrophic amoeboid flagellates that have only been identified in anoxic environments^23–26^ and possess MROs with unique FeS cluster biosynthesis (*i*.*e*., the SMS) and ETC configurations. Sparse transcriptomic data from other breviates such as *Breviata anathema* and *Lenisia limosa* showed differential retention of the SMS and ISC systems with similar patchy distributions of ETC components. Our analyses of deeply sequenced transcriptomes from diverse breviates reveal that at least three general types of MRO metabolism exist in Breviatea along a continuum of Fe-S cluster biosynthesis and ETC-related functions. By examining diverse breviates with different MRO metabolisms, we reconstruct the relative timing and evolutionary mechanisms shaping the reduction and expansion of metabolic networks within the Breviatea lineage.

## RESULTS AND DISCUSSION

### Breviatea is the deepest branch within Obazoa

To clarify the phylogenetic position and metabolic potential of Breviatea, we sequenced the transcriptomes of six breviate isolates (*Breviata anathema* and five undescribed species herein referred to as LRM1b, LRM2N6, SaaBrev, BLO, and FB10N2). Phylogenetic analysis of the SSU rRNA gene of some of these isolates previously showed that FB10N2 is closely related *Subulatomonas tetraspora* while LRM2N6 and LRM1b form sister lineages closely related to *Pygsuia* albeit with only moderate support^26^. To robustly resolve the position of these species in the context of studied breviates^23–25^, and other eukaryotes, we performed phylogenomic analyses with representatives from major lineages of eukaryotes. Consistent with previous investigations^23^, we recovered the monophyly of the supergroup Obazoa, containing Breviatea, Apusomonadida and Opisthokonta, with full branch support (Fig. 1A). Within Obazoa, the eight breviate species emerged as a well-supported sister clade to Apusomonadida and Opisthokonta in all analyses. *Pygsuia biforma*, LRM2N6, LRM1b, and SaaBrev (PLLS) formed a fully supported clade deeply nested within Breviatea with BLO as the closest sister lineage followed by *Lenisia limosa. Breviata anathema* and FB10N2 branch together with full support as sister to all other breviates (Fig. 1A, Supplementary Fig. 1).

**FIGURE 1.**
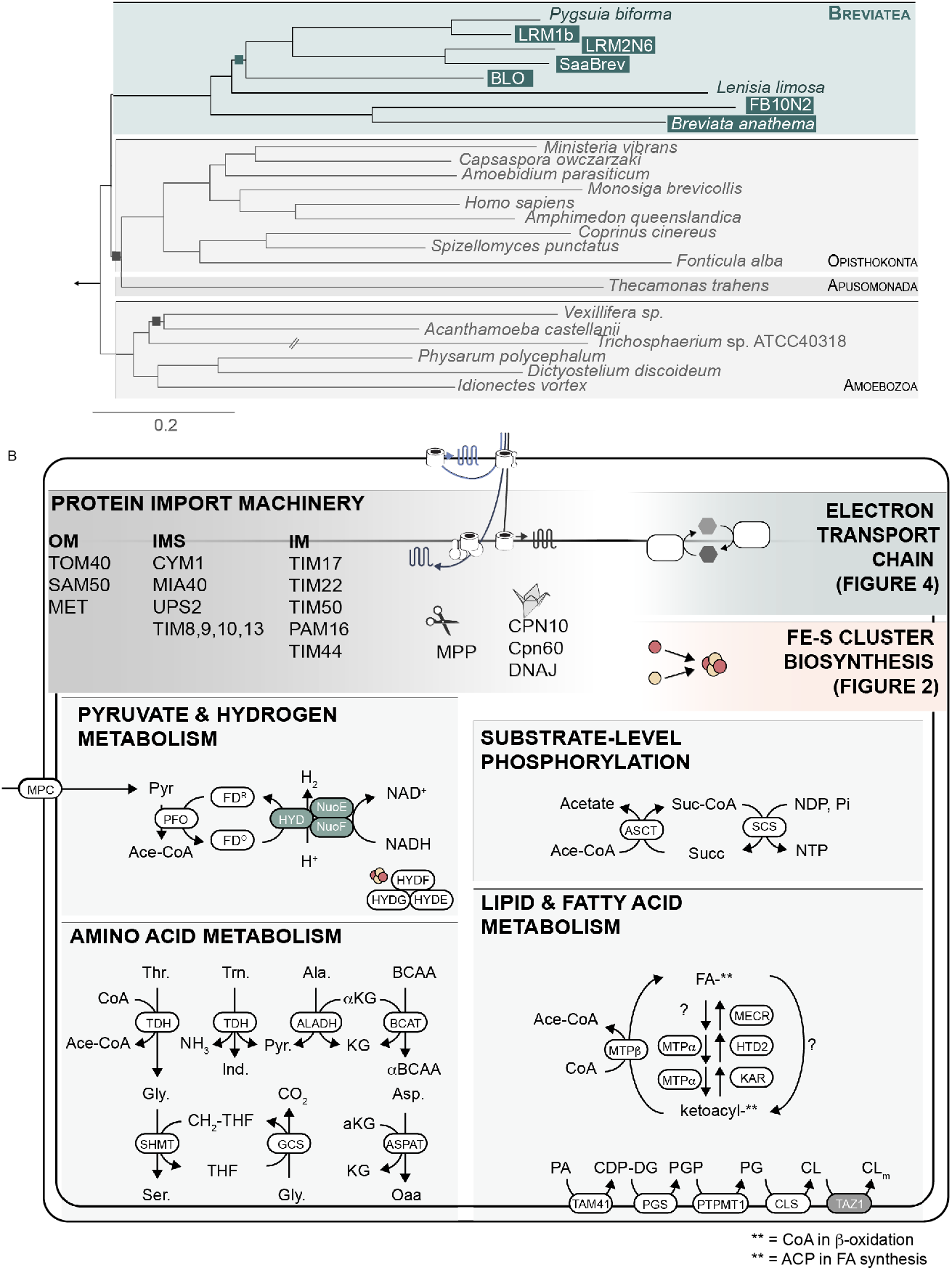
A. Breviatea representatives emerge as a clade sister to Opisthokonta and Apusomonada. Maximum likelihood phylogeny of breviate species inferred using the LG+C60+F+G model from a 253 concatenated protein phylogenomic dataset. Support values for branches were calculated using the SH-aLRT method, Bayesian posterior probabilities (PP) and nonparametric bootstrap (BS) support. All unlabeled bipartitions are supported with maximum support while squares represent bipartitions with >90% support for one or more of the computed metrics. Other eukaryotic groups were omitted for illustrative purposes; the full phylogeny can be found in Supplementary File 1 **B**. A summary of predicted MRO metabolisms conserved in most of the breviate species examined. The shaded Tafazzin1 (TAZ1) protein was only identified in *Lenisia limosa* and shaded HYD, NUOE, and NUOF were not identified in BLO. Detailed information of the predicted proteins in all pathways is listed in Supplementary File 1.

### Breviate MROs with different degrees of metabolic reduction

Previous microscopic investigations and/or *in silico* metabolic reconstructions demonstrated that the breviates *P. biforma, L. limosa*, and *B. anathema* possess MROs with different morphologies^23,24^ and predicted metabolisms^15,27^ . To compare metabolic functions across this major lineage, we predicted the MRO proteomes from the transcriptome data for each breviate species based on the detection of N-terminal mitochondrial targeting sequences (MTS) or similarity to known mitochondrial proteins^15,18^. We identified 69-107 proteins, depending on the species, that were predicted to function in the mitochondrial matrix (MM), inner membrane (IM), intermembrane space (IMS), or outer membrane (OM) (Fig. 1B, Supplementary File 1). Additionally, we identified 5-12 cytosolic proteins related to Fe-S cluster biosynthesis or pyruvate metabolism (Supplementary File 1).

The MRO proteins are predicted to function in amino acid metabolism, metabolite import and export, fatty acid metabolism, Fe-S cluster biosynthesis, lipid metabolism, lipoate metabolism, oxidative stress response, protein import and folding, pyruvate metabolism, and ATP generation (Fig. 1B). Consistent with previously described MROs, we failed to detect transcripts encoding most proteins of the ETC, tricarboxylic acid cycle (TCA), and organellar DNA maintenance and expression machinery (e.g., transcription, translation, replication). Notably, some breviate lineages have undergone more metabolic reduction than others, therefore providing a useful case study for understanding how organelles evolve. Below we discuss key pathways that have different representation within the various lineages and use these data to infer how mitochondrial metabolism has evolved in Breviatea.

### Gene gain and loss events have shaped Fe-S cluster biosynthesis in Breviatea

There are at least five pathways thought to be critical for the biosynthesis of Fe-S clusters in various compartments of eukaryotes: the mitochondrial/MRO ISC, SMS, and NIF-like systems; the cytoplasmic CIA and bacterial-like SUF system; and the plastidial SUF system^28^. Previous work^15^ showed that *P. biforma* does not encode the core ISC system components but instead encodes two isoforms of a protein that has the SMSC and SMSB subunits fused together (herein SMSCB); one of the isoforms is predicted to function in the MROs (mSMSCB) and one in the cytoplasm (cSMSCB). In contrast, core ISC components were found in the *B. anathema* data available at the time^15^. To determine whether the patchy distribution patterns of Fe-S cluster biosynthesis pathways is widespread in Breviatea, we catalogued the presence/absence of SMS and ISC in the transcriptome data from eight breviates.

#### ISC System

The mitochondrial ISC system is responsible for the maturation of 2Fe-2S and 4Fe-4S clusters in the early and late ISC pathway, respectively^29^. In the early ISC pathway of yeast, 2Fe-2S clusters are assembled on the scaffold protein ISCU where sulfur, iron, and electrons are donated by cysteine desulfurase (ISCS), frataxin (YFH1), and adrenodoxin (YAH1), respectively. This assembly is aided by accessory and chaperone proteins (ISD11, JAC1, SSQ1, and MGE1). The 2Fe-2S clusters are added to recipient proteins or further matured into 4Fe-4S proteins by the actions of IBA57, ISCA1/2, BOLA, GRX5, IND1, and NFU1.

We observed three different complements of ISC components across breviates (Fig. 2). *Lenisia limosa* encoded the most complete ISC system that notably includes early ISC components YFH1, and JAC1 and late ISC components BOLA and ISCA (Isa-like). Meanwhile BLO, FB10N2, and *B. anathema* encoded a more reduced complement, including only the core early (ISCU, ISCS) and late (NFU1, IND1) components. Finally, Saa, LRM2N6, *P. biforma* and LRM1b lacked the majority of the ISC system and have retained only the chaperone proteins (SSQ1, MGE1, NFU1, IND1). We did not detect ISD11, GRX5 or IBA57 in any of the breviates.

**FIGURE 2.**
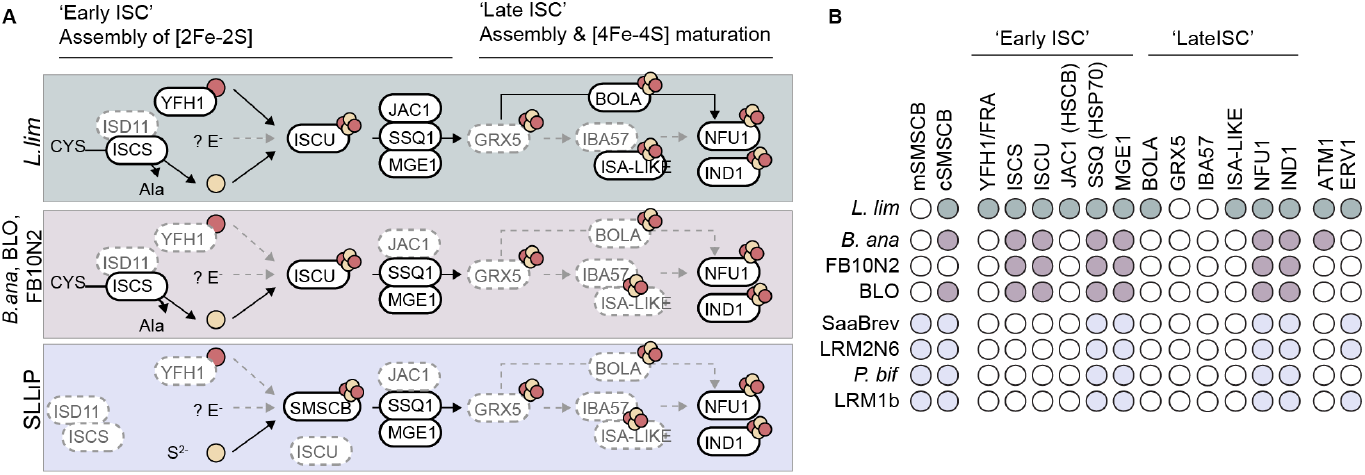
Fe-S cluster biosynthesis evolution across. **A**. Three distinct complements of FeS cluster biosynthesis machinery were detected across the indicated breviates shaded in green, pink and purple respectively. In each panel, detected and undetected proteins are outlined in solid or dashed lines, respectively. B. Presence and absence of MRO proteins are labeled in colours and blank, respectively. L. lim: *Lenisia limosa*; B. ana: *Breviata anathema*; P. bif: *Pygsuia biforma*; Saa: SaaBrev; SLL1P: SaaBrev, LRM1b, LRM2N6, and *Pygsuia biforma*.

Although ISD11 is important for the activity of ISCS in other eukaryotes, some protists do not encode clearly identifiable ISD11 orthologs, indicating that this protein may not be essential for Fe-S cluster biosynthesis or that it is difficult to detect with current methods^30–33^. The early ISC pathway relies on the chaperone complex, formed by chaperones SSQ1 and JAC1, and the ADP-ATP exchange factor MGE1^34^. However, given the consistent absence of JAC1 in most breviate MROs, we suspect that SSQ1 and MGE1 alone are sufficient for Fe-S cluster maturation. This parallels observations in other MROs where JAC1, and not SSQ1, is found to co-occur with MGE1 (*e*.*g*., in *Stygiella incarcerata*^16^). It is possible that either SSQ1 or JAC1 can fulfill the chaperone function in MRO Fe-S cluster biosynthesis.

In model eukaryotes, the late ISC pathway assembles 4Fe-4S respiratory chain components and the TCA cycle enzyme aconitase^35^ in mitochondria. The assembly of these clusters are coordinated by ISCA1/2, IBA57, and GRX5 and are ultimately added to their recipient proteins via IND1 or NFU1. Some late ISC components (e.g., ISCA1/2, BOLA, GRX5 and NFU1) have been identified in highly reduced MROs that do not possess obvious recipient proteins^16,36,37^. Therefore the absence of the late ISC pathway in most breviates is unusual as even the dramatically reduced MROs, *e*.*g.,Giardia* mitosomes, retain homologues of at least ISCA1/2, BOLA, and/or GRX5^37^. However, we cannot exclude the possibility that these genes have not been identified in most breviates due to missing data.

#### SMS Pathway

In all breviates except FB10N2, we have identified transcripts encoding putatively cytoplasmic SMSCB proteins (*i*.*e*., these proteins lack MTSs). We were also able to identify MTS-possessing mSMSCB proteins in LRM1b, LRM2N6, and SaaBrev (Fig. 2; Supplementary File 1). This large sample size provides an opportunity to investigate how the SMS pathway evolves in mitochondria. The bioinformatic analyses presented here strongly suggest that breviate species contain either a mSMSCB protein (*P. biforma*, LRM1b, LRM2N6, and SaaBre) or ISCU and ISCS (*B. anathema*, FB10N2, BLO, and *L. limosa*) in their MRO; meanwhile, the accessory components (SSQ1, MGE1, NFU1 and IND1) are conserved across the lineage. In phylogenetic analyses (Fig. 3, Supplementary Fig. 2), all eukaryotic SMSCB proteins, from breviates, stramenopiles, and the discobid *Stygiella*, form a clade that is sister to the Methanomicrobiales archaea. This suggests that genes encoding SMSCB were acquired by one of these eukaryotic lineages from a Methanomicrobiales donor^15,16,18^. The mSMSCB from the PLLS breviates form a strongly supported clade nested within the cSMSCB proteins from all the studied breviates (Fig. 3).

**FIGURE 3.**
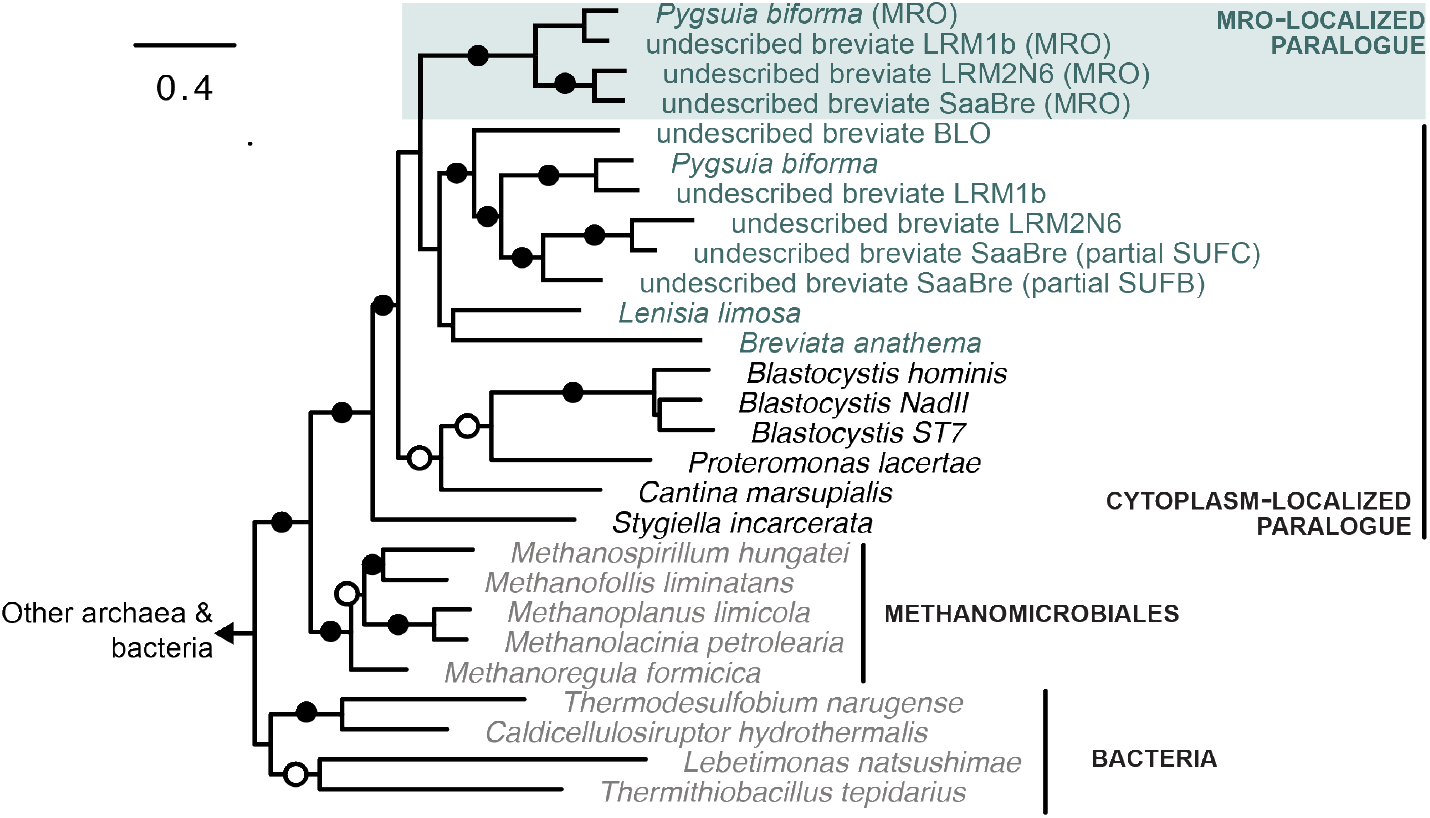
Maximum likelihood amino acid phylogeny constructed based on concatenated SMSC and SMSB homologs under the LG+C60+F+G model. Bootstrap values of ML tree are labeled near the branches. Scale bar corresponds to 0.4 substitutions per site. Tip labels are coloured in green (breviates), black (other eukaryotes) and grey (prokaryotes). Predicted sub-cellular localization for the eukaryotic proteins are indicated.

The SMS and ISC systems differ fundamentally in their mode of action: the ISC system relies on cysteine-derived sulfur liberated by ISCS, while the archaeal SMS system utilizes environmental sulfide^38^ (a common molecule in hypoxic marine sediments). Moreover, the ISC system depends on iron delivered by frataxin ^34^ while the SMS system utilizes an unknown iron source and might use intracellular Fe(II)^14,39^. Despite these differences, the two systems perform similar roles. For example, it was shown that the SMS proteins from *Methanocaldococcus jannaschii*^40,41^ were capable of complementing *E. coli* mutants lacking the SUF and ISC systems in the presence of sulfide^40,41^. This suggests that the SMS proteins can perform analogous roles to the SUF and ISC systems.

We propose that the ancestor of Breviatea possessed a canonical mitochondrial ISC pathway and a laterally acquired a SMSCB gene whose product initially functioned in the breviate cytoplasm (‘SMSCB’ Fig. 5). The ISC system and cSMSCB were maintained by various lineages including *Breviata, Lenisia* and BLO, with secondary loss in FB10N2 (although we cannot exclude its absence because of incomplete transcriptome coverage). There was a gene duplication of *smscb* in the lineage that gave rise to the PLLS clade and one of the copies acquired a mitochondrial signal sequence (mSMSCB). The mSMSCB was likely functionally redundant with the existing ISC system permitting the eventual loss of the latter in the PLLS clade.

#### Mitochondrial-cytoplasm crosstalk

Previous studies suggest that an unknown factor ‘X’ produced by the ISC system is exported to the cytosol by ATM1, an ABC transporter in the mitochondrial inner membrane^29,33,35^. Of the eight breviates, we were only able to identify an ATM1 in *L. limosa* and *B. anathema* (Supplementary File 1; Fig. 2). ERV1, a sulfhydryl oxidase located in the intermembrane space, which is required for the export of ‘factor X’, was only detected in LRM1b and LRM2N6 (Supplementary File 1; Fig. 2). These components show variable patterns of retention and loss across anaerobic protist lineages^16,42^. Since our analyses are based on transcriptomes, it is not possible to distinguish whether the lack of the detection of these genes above indicates they are truly absent or simply the result of incomplete transcriptome coverage.

### Breviate MROs likely couple hydrogen production to substrate-level phosphorylation

#### Pyruvate, hydrogen and energy metabolism

In the mitochondria of aerobic eukaryotes, the pyruvate dehydrogenase complex (PDC) oxidizes pyruvate to acetyl-CoA which enters the TCA cycle to ultimately generate reducing equivalents in the form of NADH and succinate. These reducing equivalents are fed into the ETC which generates a proton gradient that fuels ATP generation by oxidative phosphorylation coupled to the reduction of oxygen. Pyruvate metabolism is generally different in MROs of anaerobic protists, including breviates. Some anaerobes do not encode a PDC but instead rely on an MRO-localized pyruvate:ferredoxin oxidoreductase (PFO) that is predicted to oxidize pyruvate to acetyl-CoA with the concomitant production of reduced ferredoxin and CO_2_. The ferredoxin is reoxidized and electrons transferred to protons by the action of an [FeFe]-hydrogenase (HYDA) to yield hydrogen gas^4^. We detected transcripts encoding at least one putatively MRO-localized PFO and one ferredoxin protein in all species of breviates. Our result suggests there were three PFO copies present before the divergences of all these eight breviate species (Supplementary Fig. 3).

We also identified genes encoding the pyruvate carriers MPC1 and MPC2 in *Pygsuia*, LRM1b, *L. limosa* and BLO, that may be responsible for pyruvate import into MROs in these species. All breviates encoded MRO-targeted hydrogenase maturase proteins, indicating that the machinery for hydrogenase maturation within MROs is conserved across the lineage. In contrast, while most breviates encoded an MRO-targeted HYDA, BLO lacked a clearly identifiable MRO-localized HYDA. Instead, BLO encoded HYDA homologues that appear to be cytoplasmic based on their domain composition and lack of MTS. Whether the absence of an MRO-targeted HYDA in BLO is the result of incomplete data, cryptic organelle targeting, or bona fide absence cannot be determined with present data; however, the absence of other HYDA-associated proteins (see NUOE, NUOF discussion below) suggests that BLO does not express a mitochondrial HYDA under the culturing conditions employed in this study.

The acetyl-CoA from the PFO reaction is predicted to be converted to acetate by acetate:succinate CoA transferase (ASCT) and succinyl-CoA synthetase (SCS) to generate ATP by substrate-level phosphorylation. We identified transcripts predicted to encode MRO-localized ASCT 1B, ASCT 1C and SCS subunits in all species of breviates. Previous cell biological investigations of *P. biforma* demonstrated that antibodies raised against ASCT colocalize with mitochondrial dyes, strongly suggesting that this pathway occurs in the MRO^15^. In summary, our results suggest that all studied breviate species produce ATP by the anaerobic ‘hydrogenosomal’ pathway that includes ASCT and SCS.

#### TCA Cycle

In aerobic eukaryotes, acetyl-CoA is oxidized by the enzymes of the TCA cycle, generating reducing equivalents such as NADH that are essential for the ETC. In MROs, the TCA cycle and ETC are often highly reduced or lost completely. We failed to detect transcripts encoding aconitase, isocitrate dehydrogenase, and α-ketoglutarate dehydrogenase in any of the breviates; neither did we find citrate synthase in LRM2N6, SaaBrev, *L. limosa*, BLO, *B. anathema*, and FB10N2 (Supplementary File 1). We also observed a variable pattern of succinate dehydrogenase (SDH), and fumarate hydratase (FH) in the breviate lineage (Supplementary File 1, Fig. 4A, B). These findings suggest that the TCA cycle is incomplete and that individual components might perform the reverse reactions or participate in other pathways as previously reported^3^.

**FIGURE 4.**
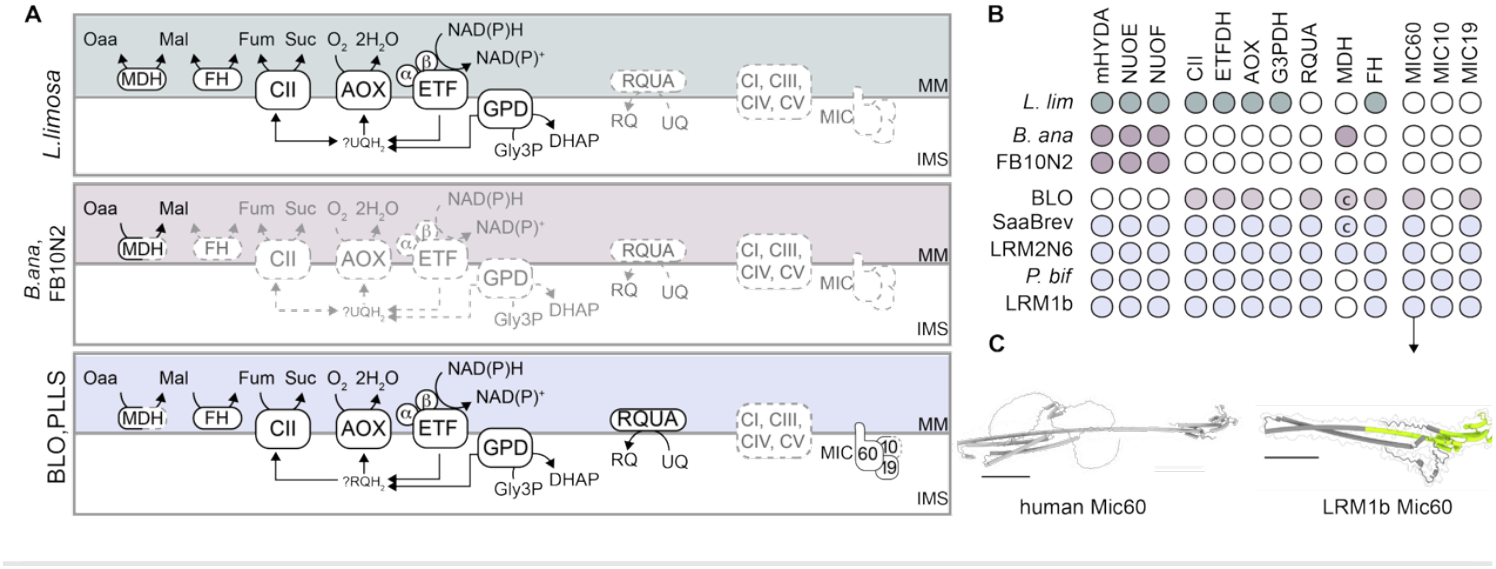
ETC and MICOS components are differentially retained across Breviatea. **A**. Three main categories of ETC and MICOS complements were detected across the indicated breviates shaded in green, pink and purple respectively. In each panel, detected and undetected proteins are outlined in solid or dashed lines, respectively. Proteins detected in only some of the indicated representatives are shown in partially dashed outlines. **B**. Presence and absence of ETC and MICOS components are shaded or unshaded, respectively. Proteins predicted to function in the cytoplasm are indicated with ‘c’. **C**. Structure of human Mic60 (left) and AlphaFold-resolved of LRM1b Mic60 (right). *L. lim*: *Lenisia limosa*; B. ana: *Breviata anathema*; P. bif: *Pygsuia biforma*; Saa: SaaBrev; SLL1P: SaaBrev, LRM1b, LRM2N6, and *Pygsuia biforma*.

We did not find a malate dehydrogenase (MDH) closely related to the mitochondrial paralogue in any of the breviates. Instead, SaaBrev and LRM2N6 appear to have bacteria-like MDH homologues that possess an N-terminal extension and putative MTS. In phylogenetic reconstructions, the SaaBrev and LRM2N6 MDH sequences branch together in a clade formed by sequences from Bacillati with weak support (BP_uf_ = 72), suggesting that this gene was likely acquired through lateral transfer whose donor cannot be determined with present data (Supplementary Fig. 4). Nevertheless, we propose that SaaBrev and LRM2N6 import malate into the MRO using the malate/aspartate shuttle and convert it to oxaloacetate via MDH.

### Some breviates possess a truncated ETC and cristae organization complex

We did not identify any transcripts encoding components of CIII, CIV, or CV in any of the breviates. All breviates except BLO encode two subunits of CI (NUOE and NUOF). The retention of only the NUOE and NUOF subunits of CI has been reported in other MROs, including the hydrogenosomes of *Trichomonas vaginalis*^43^. It is thought that these soluble subunits do not function in a membrane-associated ETC but instead coordinate with [FeFe]-hydrogenase to couple NADH oxidation to hydrogen production, as observed in some anaerobic prokaryotes^44^. The absence of detectable NUOE and NUOF-encoding transcripts in BLO correlates with their lack of a putative MRO-localized HYDA suggesting NADH-dependent hydrogen production does not occur in the organelle.

All breviates except *B. anathema* and FB10N2 encode CII and quinone-utilizing complexes such as electron transferring flavoprotein dehydrogenase (ETF, ETFDH), alternative oxidase (AOX) and glycerol-3-phosphate dehydrogenase (G3PDH) (Supplementary File 1, Fig. 4). We detected transcripts encoding the RQ-biosynthesis protein RQUA in the PLLS group and in BLO, but not in *L. limosa, B. anathema* and FB10N2 (Fig. 4). This suggests that quinone-dependent electron transport chains were likely lost in MROs of the *B. anathema* and FB10N2 lineage (Fig. 5). The lack of obvious RQ-biosynthesis machinery in *L. limosa* could be due to missing data or could indicate that RQUA was acquired by breviates after the split between *L. limosa* and the BLO+PLLS grouping.

**FIGURE 5.**
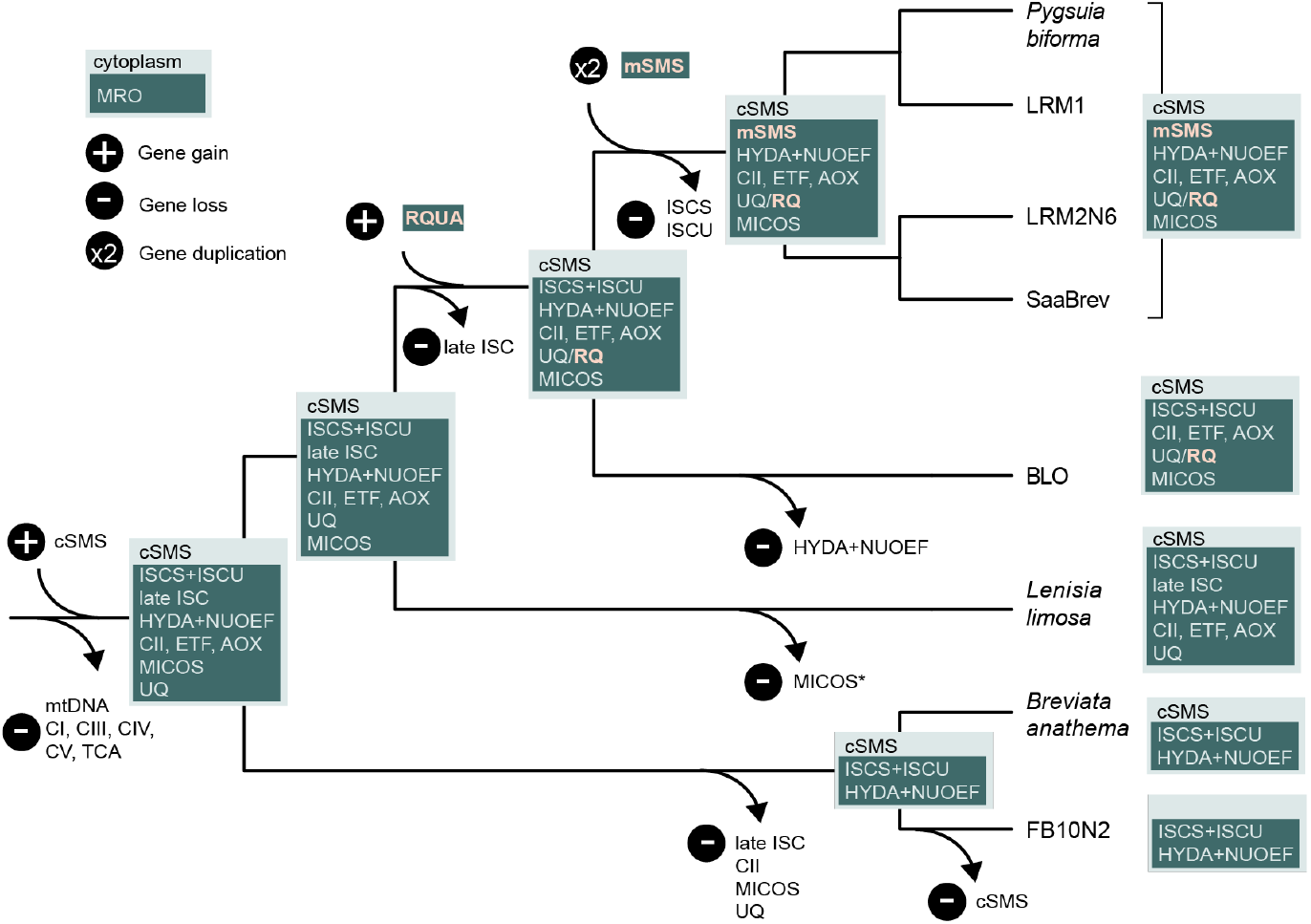
A Hypothesis of the Evolutionary history of MRO metabolism in breviates. Hypothesized ancestors are depicted at the nodes as inferred from modern representatives (right hand schematics). Predicted MRO-localized proteins are labeled in dark green boxes. Features acquired after the evolution of the last common ancestor of breviates sampled herein are shown in orange.

In aerobic eukaryotes, the mitochondrial inner membrane extends through invaginations known as cristae, which are important for submitochondrial organization of the respiratory chain. The mitochondrial contact site and cristae organizing system (MICOS) is a multi-subunit protein complex that forms and maintains mitochondrial cristae^45^ and consists of core (MIC60, MIC10 and MIC19)^46^ and accessory (MIC12, MIC25, MIC26 and MIC27) subunits^47,48^. Although highly conserved among aerobic eukaryotes, the core MICOS subunits are patchily distributed or completely absent in anaerobic lineages^46^. This could be a consequence of having a reduced (or absent) ETC, thereby relaxing the selection for the retention of cristae maintenance proteins. To date, MICOS has not been reported in Breviatea^46^, and it remains unresolved whether cristae exist within the breviate clade, with only a single report of tubular cristae in *B. anathema*^49^. Homologues of MIC60 and MIC19 were identified in SaaBrev, LRM1b, LRM2N6, *P. biforma* and BLO while MIC10 candidates were detected in only in LRM1b and *P. biforma* (Fig. 4B). The predicted MIC60 structures from the breviates adopt the Mitofilin fold characteristic of MIC60 (Fig. 4C). Phylogenetic reconstruction of MIC60 homologues across Obazoa showed that all Breviatea sequences form a well-supported, clade sister to metazoa and unicellular holozoa (Supplementary Fig. 5). However, we failed to recover monophyletic Opisthokonta likely owing to the extremely divergent nature of the yeast homologues that has previously been reported^50^.

We could only confidently detect MICOS components in breviates that encode Q-utilizing complexes and the RQ-biosynthesis protein RQUA (Fig. 4, 5). The absence of MICOS in *B. anathema* and FB10N2 is unsurprising, given limited ETC components that remain in the MRO (Fig. 4). Interestingly, no MICOS proteins could be detected in the transcriptome or genome *L. limosa*, the only other breviate that encodes Q-utilizing complexes. This could be due to missing data, as both the transcriptome and genome are highly fragmented, or represent a true absence. The co-occurrence of MICOS and CII may reflect a minimal level of ETC activity associated with cristae maintenance, but the predicted absence of MICOS in *L. limosa* would indicate this relationship is not mandatory. Truncated respiratory chains have been identified in some anaerobic ciliates^51,52^, rhizarians^53^, stramenopiles^54^, amoebozoans^55,56^ and chytrid fungi^21,57^ some of which retain MICOS components^46^. In all cases, CII and at least one other Q-utilizing component is conserved demonstrating that Q-based electron transfer and a minimal MICOS complex can be retained independent of oxidative phosphorylation. The identification of MICOS components suggests that some breviates might possess small cristae that have evaded detection to date, although only a minority of the taxa discussed here have been examined using electron microscopy. Alternatively, it could indicate that MICOS components have additional functions other than cristae maintenance, perhaps related to redox balance, protein import, or organelle division as reported for some MICOS^48^. Therefore, functional interrogation of the minimal breviate MICOS complex in these cristae-lacking MROs could, in the future, uncover alternative functions that have been overlooked in model system mitochondria.

## CONCLUSIONS

The evolution of mitochondrion-related organelles in the breviate lineage appears to involve the interplay of lateral gene transfer, gene duplication, subcellular retargeting, and gene loss, especially within the Fe-S cluster biosynthesis pathways and the ETC. The last common ancestor of the breviates investigated herein possessed a mitochondrial ISC pathway and a SMSCB protein that functioned in the cytoplasm. The gene duplication of SMSCB and relocalization of the mSMSCB to the MRO of the ancestral lineage of the PLLS breviates led to functional redundancy with the existing ISC system, thereby allowing the eventual loss of the mitochondrial ISC system in the PLLS lineage. Interestingly, the variation in Fe-S cluster biosynthesis machinery components over breviate diversity is not clearly correlated with the presence/absence of MRO proteins or cytoplasmic Fe-S cluster biosynthesis proteins. We observed different evolutionary trajectories with respect to the respiratory chain where some lineages have completely lost all quinone-metabolism and cristae organizing proteins while others retain a streamlined respiratory chain and core MICOS proteins. Overall, we observed distinct trajectories for different components of the MRO proteomes over the breviate lineages (Fig. 5). This is an interesting example of the macroevolutionary concept of “mosaic evolution”; *i*.*e*., the evolution of different structures of an organism at different rates^58^. Here, evolutionary changes occur to different biochemical modules in the MRO in different lineages in adaptation to similar low oxygen and sulfidic environments with differential retention of ancestral features in each lineage. Determining whether or not this is of adaptive significance to the different MRO types will require further investigation. However, we suspect that the availability of environmental sulfide was important for the fixation of the sulfide-dependent SMS system. It is possible that at least some of the differences we observed relate to the different availabilities of nutrients or metabolites in the specific microbial communities and environments in which these organisms are found.

## MATERIALS AND METHODS

### Sampling, RNA Isolation, Sequencing

Protist isolates FB10N2 (isolated from anoxic marine intertidal mudflat sediment in False Bay, San Juan Islands, WA, United States of America, 48°29’07.9”N 123°04’31.1”W), SaaBrev (isolated from marine mud at low tide in British Columbia, Canada, 49°00’N 123°01’W), LRM1b and LRM2N6 (isolated from anoxic marine subtidal sediment near La Ràpita, Spain, 40°35’33.2”N 0°42’37.0”E) and BLO (originally CARMGS BMO; isolated from anoxic marine mangrove sediment near CARMABI Foundation, Piscaderabaai, Curaçao, 12°08’14.5”N 68°58’03.2”W), were sampled as described by Eglit ^59^ (2022). All isolates were monoprotistan but contained diverse prokaryotes (including prey) and were grown in Corning culture flasks filled with autoclaved or filtered seawater (sourced from the coast off Halifax, N.S., Canada), supplemented 3% v/v with liquid Lysogeny broth (LB) medium. For each sample, RNA was isolated from 0.4-2 L cultures 10-14 d after transferring, using TRIzol reagent (Invitrogen, Life Technologies, USA) or PureLink RNA Mini Kit (Thermo Fisher, USA) according to the manufacturer’s instructions. Preparation of Illumina libraries and a poly-A tail purification step to enrich for eukaryotic messenger RNA (mRNA) were performed by Genome Quebec (Canada).

### Transcriptome Assembly and Decontamination

Raw sequencing data was deposited into GenBank under accession numbers ERR14889641-ERR14889646. Adapters and low quality sequences were removed using Trimmomatic v0.36^60^, and then assembled using Trinity v2.4.0 ^61^ with default parameters. WinstonCleaner (https://github.com/kolecko007/WinstonCleaner) was used to eliminate cross-contamination from other eukaryotes in the same sequencing lane. Plast v2.3.2^62^ was used to do remove sequences similar to prokaryotes. After decontamination, 27,833 (*B. anathema*), 34,146 (BLO), 37,226 (FB10N2), 37,258 (SaaBre), 50,454 (LRM1b) to 75,410 (LRM2N6) transcripts were used for downstream analyses.

### Detection of Nucleus-Encoded MRO Proteins

Putative MRO proteins were detected by blastp searches, using a seed protein database as queries, which contained predicted or experimentally determined mitochondrial proteomes of breviate *Pygsuia biforma*^15^, *Homo sapiens* (MitoCarta2.0:https://www.broadinstitute.org/files/shared/metabolism/mitocarta/human.mitocarta2.0.html) and *Saccharomyces cerevisiae*^63^, as well as the main components of mitochondrial metabolic pathways of other anaerobic protists such as *Paratrimastix pyriformis*^20^, *Mastigamoeba balamuthi*^10^, and *Brevimastigomonas motovehiculus*^53^. Each candidate protein identified in this way was used as a query in a blastp search against the NCBI nr database to retrieve potential homologs for phylogenetic analyses. Sequences were aligned using MUSCLE trimmed using BMGE^64^ with default settings and then phylogenetic trees were constructed using IQ-TREE v 1.6.6^65^. Mitochondrial-targeting signal (MTS) prediction was performed using multiple software programmes, including Mitoprot ^66^, MitoFates^67^, TargetP^68^, and NommPred^69^.

### Phylogenomic Analyses

A 253-protein phylogenomic dataset including protein orthologs from main eukaryotic supergroups and all breviate species was constructed as previously described^70^. Briefly, orthologues of newly added species were screened against seed datasets using tblastn or blastp, and the top five hits with e-value < 10^-10^ were retained. Preliminary single-gene trees were aligned using MAFFT v7.310^71^, and then ambiguously aligned positions were trimmed using BMGE using default settings^64^. Single trees constructed by FastTree v1.0.1^72^ were manually inspected, and paralogues were removed. A final concatenated alignment with 63 taxa and 67,000 amino acid positions was used for following phylogenomic analyses. A maximum-likelihood (ML) tree was constructed by IQ-TREE v 1.6.6^65^ under the LG+C60+F+G model. Support values were calculated using the SH-aLRT support and Posterior Mean Site Frequency bootstrap support (100 replicates).

### Detection and phylogenetic analysis of MICOS proteins

Sequences of MIC60, MIC10, MIC19, and MIC26 from representative Amorphea organisms were retrieved from UniProt or NCBI and used as queries to search the generated breviate transcriptomes using TBLASTN (E-value < 1.0) (Supplementary Data). Candidate hits were translated into the appropriate reading frame and annotated InterPro^73^ and the best scoring BLASTP against the clustered NR database (date accessed: September 2025), and Foldseek (Release 10-941cd33)^74^ against the AlphaFold3-predicted structure database (AFDB50 (AlphaFold/UniProt50 v6))^75^. Newly identified breviate MICOS homologs were subsequently used as additional queries in iterative rounds of searching using similar methods to confirm their absence or presence in the other breviates. Phylosuite v2^76^ was used to conduct, manage and streamline the analyses. Sequences were aligned with MAFFT v7.505^71^ using ‘L-INS-i (accurate)’ strategy and normal alignment mode. Gap sites were removed with trimAl v1.2rev57^77^ using “-gappyout” command. The maximum likelihood tree was constructed using IQ-TREE v under the LG+C60+F+G model with 1000 ultrafast bootstrap replicates and aLRT support.

### Phylogenetic Analyses of SMSCB and PFO proteins

At least one SMSCB copy was detected for each our six newly sequenced transcriptomes except for BLO. SMSCB amino acid sequences for eukaryotes as well as the SMSC amino acid sequence for *Blastocystis hominis* were downloaded from GenBank. Using the SMSB sequence of *P. biforma* as a reference, the top 1,000 related sequences from GenBank were downloaded and then compressed into 100 sequences with >60% genetic similarities using cd-hit. Subsequently, SMSC and SMSB amino acid sequences were aligned using AliView v1.23^78^, respectively. After manually trimming and deleting ambiguously-aligned sites, 249 and 414 positions were kept and combined for inferring phylogenetic trees. A Bayesian inference (BI) analysis was performed with PhyloBayes v4.1^79^ using the CAT model. 121,000 cycles were run with four chains, and the first 20% generations were discarded as burn-in. The remaining trees were used to generate a consensus tree and to calculate the posterior probabilities (PP) of all branches using a majority-rule consensus approach. The Maximum-likelihood (ML) tree was constructed using IQ-TREE v 1.6.6^65^ under the LG+C60+F+G model with 1,000 ultrafast bootstrap replicates and aLRT support. The PFO tree was constructed in the same way as the SMSCB tree.

## Supporting information

Supplementary Results and Figures

## ACKNOWLEDGMENTS

Many thanks are due to Noèlia Carrasco Querol, Greg Gavelis and Camille Poirier for help in sampling the organisms used, and Sarah Shah, Dandan Zhao, Marlena Dlutek, Dr. Jon Jerlström-Hultqvist and Viktor Torblöm, for their help on data analyses and/or culturing. This work was supported by a Discovery Grant (2017-06792) awarded to AJR from the Natural Sciences and Engineering Research Council of Canada, the Arkansas Bioscience Institute (USA) awarded to JDS, and the National Natural Science Foundation of China (grant numbers 32070406) awarded to ZY. And China Scholarship Council provided support for training ZY at Dalhousie University.

## DATA AVAILABILITY

The data underlying this article are available in the article, online supplementary material, and figshare databases. Raw sequencing data was submitted to GenBank under ERR14889641-ERR14889646.

