## Supplementary Results and Figures for "DIVERGENT TRAJECTORIES FOR ANAEROBIC MITOCHONDRIAL EVOLUTION IN BREVIATE PROTISTS"

### SUPPLEMENTARY RESULTS AND DISCUSSION

#### *The CIA pathway is partially conserved in Breviatea*

The CIA pathway located in the cytosol of eukaryotes is necessary for biosynthesis of cytosolic and nuclear Fe-S proteins, that play key roles in ribosome maturation, transcription, and DNA repair-replication [1,2]. We found that the transcriptomes of LRM2N6 and SaaBre encode proteins comprising a complete CIA pathway, while the remaining breviate transcriptomes encode only some components (Supplementary File 1). Across anaerobic protists, there is no clear pattern in terms of which CIA system components are present in lineages predicted to use a MRO SMSCB (mSMSCB), an MRO ISC system or the bacterial-like SUF pathway[3].

CFD1-NBP35 serves as scaffold for assembly of cytoplasmic 4Fe-4S clusters [2,4]. While NBP35 is universally conserved among eukaryotes, CFD1 displays a patchier distribution[5] suggesting that NBP35 might be sufficient for cytosolic Fe-S cluster assembly. Indeed, NBP35 is present in all eight breviate species, but CFD1 is not detected in *B. anathema* and FB10N2. In model systems, electrons from NADPH are donated to Fe-S cluster biosynthesis by the action of TAH18 and DRE2 (yeast) or NDOR1 and CIAPIN1 (human) [2,6]. We could not detect homologues of DRE2, NDOR1, or CIAPIN1 in any of the breviate transcriptomes. However, we did identify putative TAH18 proteins in LRM2N6, SaaBrev, and *L. limosa* which are missing in other anaerobes [3]. This indicates that other components in anaerobic protists might functionally replace TAH18-DRE2/NDOR1-CIAPIN1, or that biosynthesis of cytosolic and nuclear Fe-S proteins in anaerobic protists does not need these components.

In the last step of the CIA pathway, NAR1, CIA1, CIA2, and MMS19 transfer the 4Fe-4S cluster from the scaffold to a target apoprotein[7]. Previous work demonstrated that the core components NAR1, CIA1 and CIA2 are well conserved across the tree of eukaryotes while MMS19 has been lost numerous times[5]. Similarly, we found that most breviates except *B. anathema* encode all three core components of the late CIA pathway, while *L. limosa* and FB10N2 do not encode MMS19.

### SUPPLEMENTARY FILES

Supplementary File 1 – Spreadsheet containing predicted proteins from newly sequenced breviates.

**SUPPLEMENTARY FIGURES**

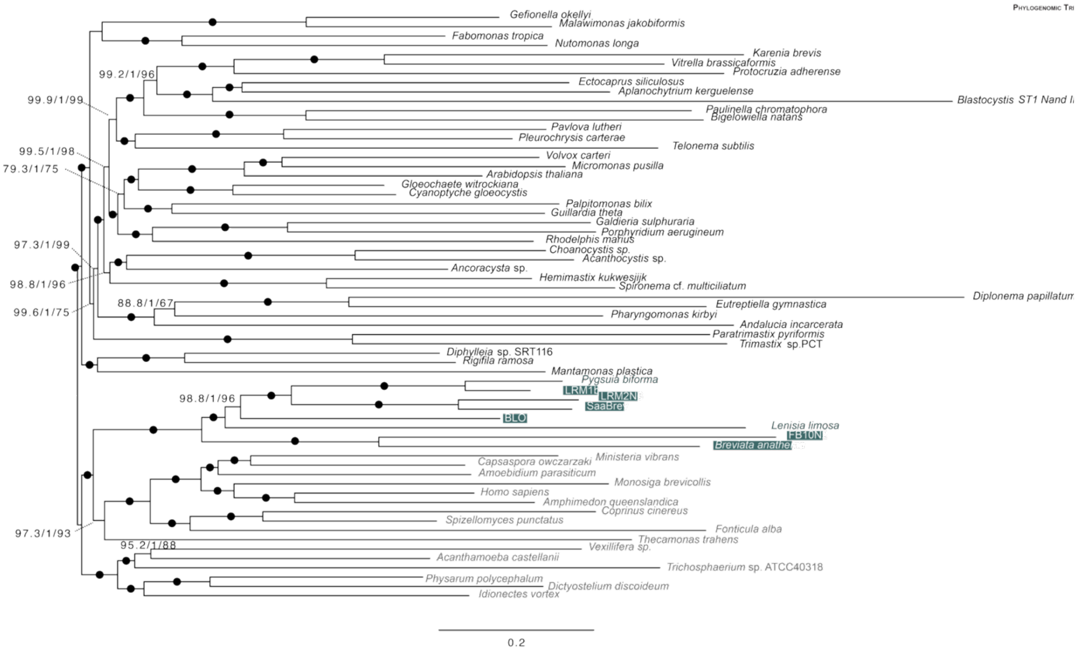


**SUPPLEMENTARY FIGURE S1** – Breviatea representatives emerge as a monophyletic clade sister to Opisthokonta and Apusomonada. Maximum likelihood phylogeny of breviate species inferred using the LG+C60+F+G model from a 253 concatenated protein phylogenomic dataset. Support values for branches were calculated using the SH-aLRT support (%) / aBayes support / standard bootstrap support. Closed circles on bipartitions maximum support.

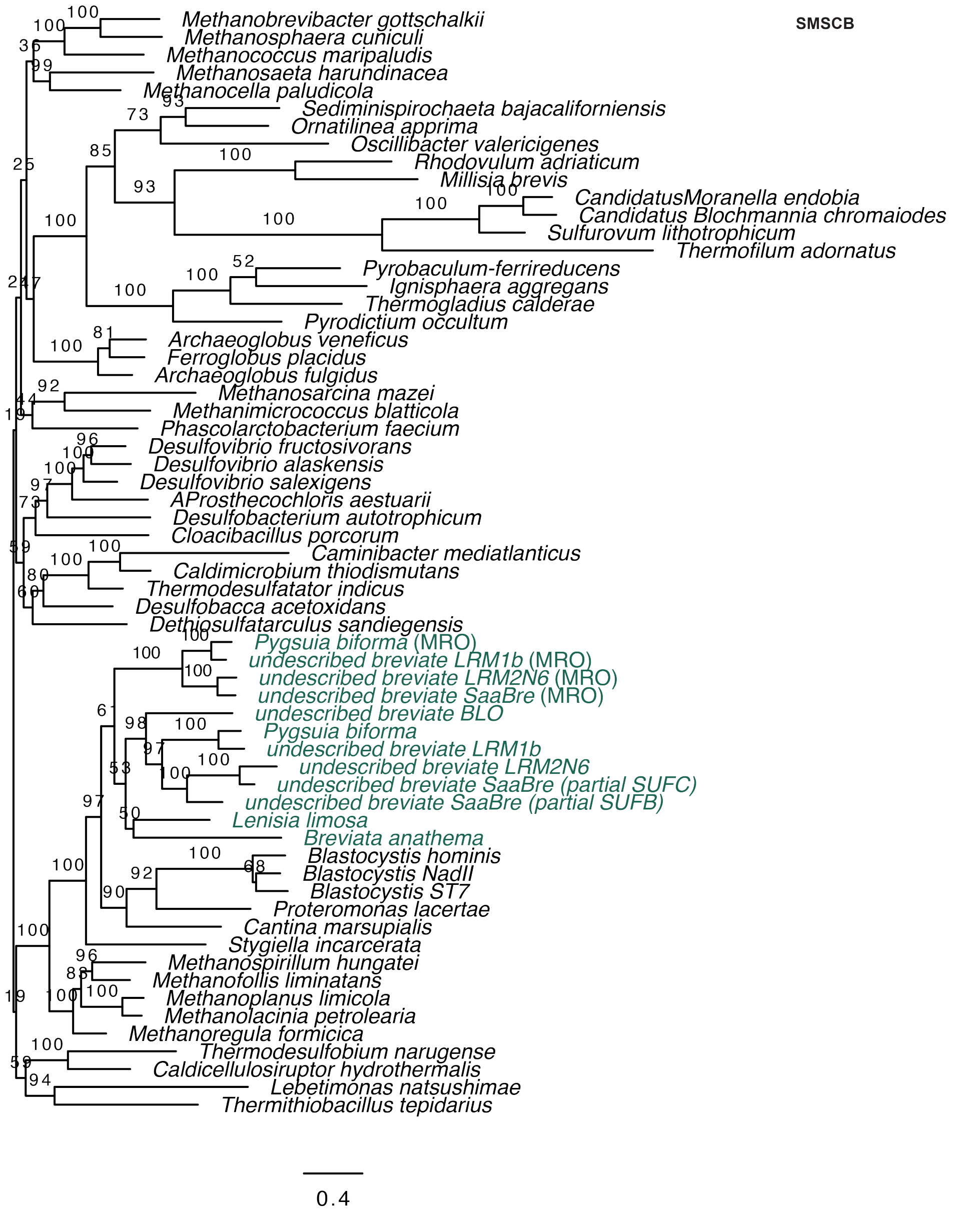


**SUPPLEMENTARY FIGURE S2 -** Maximum likelihood amino acid phylogeny constructed based on concatenated SMSC and SMSB homologs under the LG+C60+F+G model. Bootstrap values of ML tree are labeled near the branches. Scale bar corresponds to 0.4 substitutions per site. Tree file with additional support metrics can be found in the figshare respository. Tip labels are coloured in green (breviates)

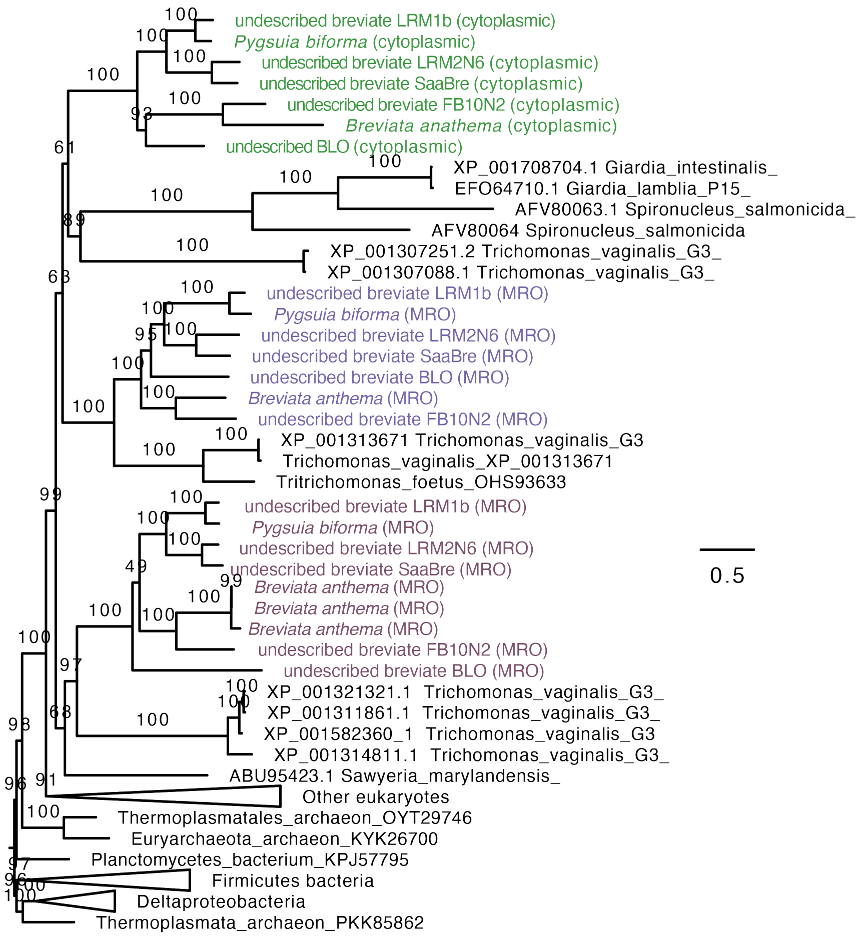


**SUPPLEMENTARY FIGURE S3 -** Maximum likelihood amino acid phylogeny of PFO homologs under the LG+C60+F+G model. Bootstrap values of ML tree are labeled near the branches. Scale bar corresponds to 0.5 substitutions per site. Three PFO paralogues from breviates are coloured in green, purple and pink.

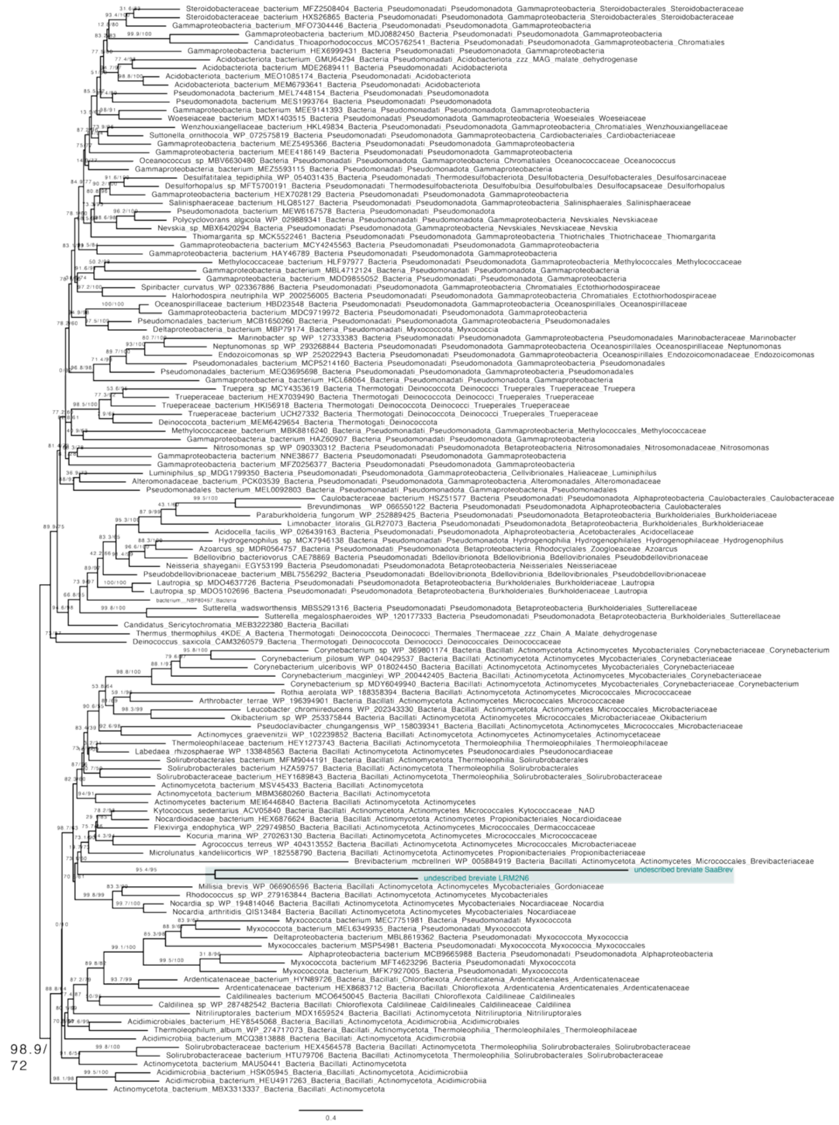


**SUPPLEMENTARY FIGURE S4 -** Maximum likelihood amino acid phylogeny constructed based on MDH holomogues under the LG+C60+F+G model. SH-aLRT and ultrafast bootstrap values are labeled near the branches. Scale bar corresponds to 0.4 substitutions per site. Tip labels are coloured in green (breviates).

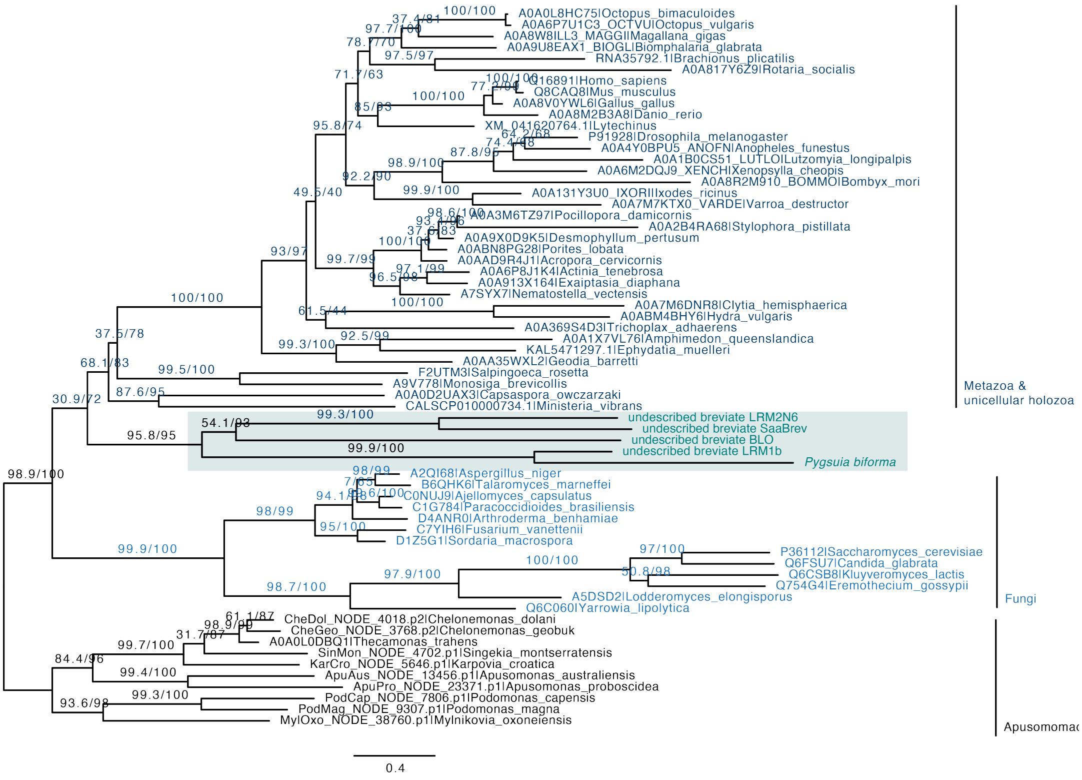


**SUPPLEMENTARY FIGURE S5 -** Maximum likelihood amino acid phylogeny of MIC60 homologs under the LG+C60+F+G model. SH-aLRT support (%) and ultrafast bootstrap values of are labeled near the branches. Scale bar corresponds to 0.4 substitutions per site. Breviates are labeled in green and Opisthokonts in different shades of blue.

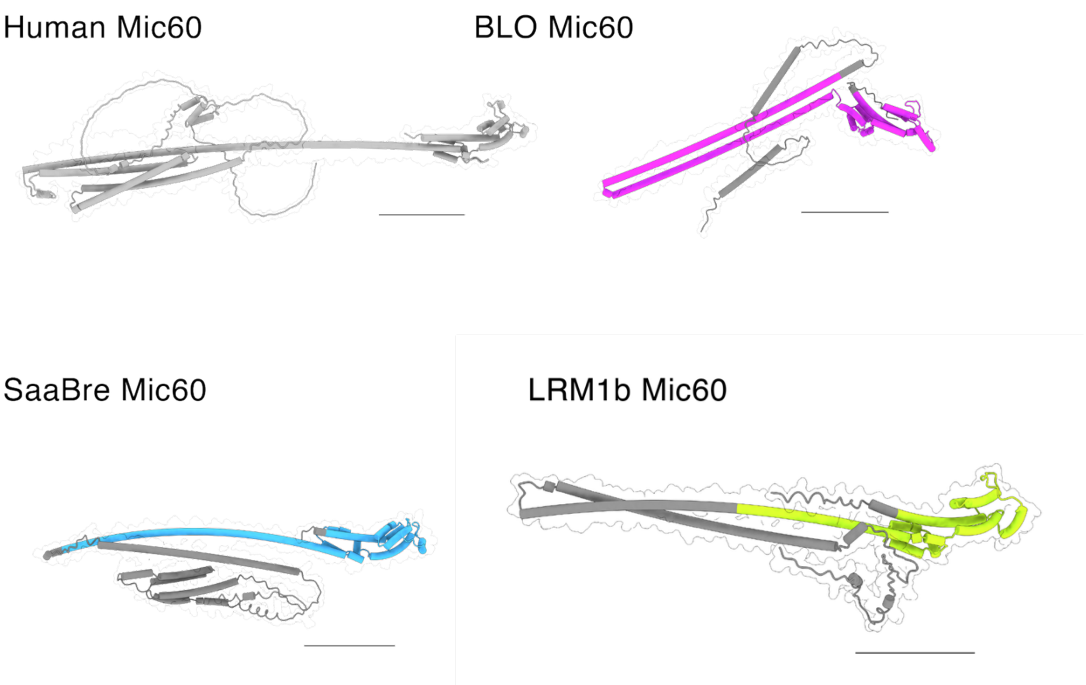


**SUPPLEMENTARY FIGURE S6** AlphaFold3 predicted Mic60 structures of human (UniProt: Q16891, grey), BLO (pink), SaaBre (blue) and LRM1b (yellow).  The mitofilin domain highlighted. Scale bar, 50 Å.

98
